# Fresh insights into Mediterranean biodiversity: Environmental DNA reveals spatio-temporal patterns of stream invertebrate communities on Sicily

**DOI:** 10.1101/2021.06.10.447858

**Authors:** Kamil Hupało, Saskia Schmidt, Till-Hendrik Macher, Martina Weiss, Florian Leese

**Affiliations:** Aquatic Ecosystem Research, Faculty of Biology, University of Duisburg□Essen, Essen, Germany; Department of Animal Ecology, U4, German Federal Institute of Hydrology, Koblenz, Germany; Centre for Water and Environmental Research (ZWU), University of Duisburg-Essen 45141 Essen, Germany

**Keywords:** eDNA, macroinvertebrates, freshwater, community, metabarcoding

## Abstract

The Mediterranean region with its islands is among top biodiversity hotspots. It houses numerous freshwater taxa with a high rate of endemism, but is heavily impacted by anthropogenic pressures and global climate change. To conserve biodiversity, reliable data on species and genetic diversity are needed especially for the scarcely known insular freshwater ecosystems. Environmental DNA metabarcoding provide a straight-forward opportunity to assess aquatic biodiversity. Therefore, we conducted the first eDNA metabarcoding study in one stream catchment on Sicily. Specifically, we aimed to i) investigate spatial diversity patterns of macroinvertebrate communities, ii) assess seasonal changes, and iii) check if dispersal barriers can be identified. Water samples were taken at 27 different sites in two seasons and eDNA metabarcoding performed using the COI gene. In total, we detected 98 macroinvertebrate species, including 28 taxa potentially new to Sicily. Exact sequence variant (ESV) and species composition data showed that diversity differed between seasons with less taxa detected in winter. We also detected a dispersal barrier, which had a stronger effect in autumn. Our findings show that eDNA metabarcoding provides valuable information on Sicilian freshwater biodiversity. We therefore encourage its application for understudied regions to better understand the state and dynamics of freshwater biodiversity.

## Introduction

Freshwater ecosystems are hotspots of biodiversity (Strayer and Dudgeon 2010). Even though they only comprise roughly 1% of global land surface area (Dudgeon et al. 2006; Strayer and Dudgeon 2010), freshwater ecosystems host nearly 9.5% of Earth’s described animal species (Balian et al. 2007). However, degradation of freshwater ecosystems is a global phenomenon, affecting in particular riverine habitats (Grill et al. 2019; Vörösmarty et al. 2010). Anthropogenic pressures are superimposed by the effects of global climate change, in particular heat waves, floods and droughts (Arnell 1999; Vörösmarty et al. 2010). As a consequence, freshwater faunal biodiversity is declining, making rivers and lakes the most endangered ecosystems in the world (Almond et al. 2020; Dudgeon et al. 2006; Reid et al. 2019). This holds true in particular for the Mediterranean region (De Figueroa et al. 2013), which is considered as one of the top biodiversity hotspots in the world (Médail and Quézel 1999; Myers et al. 2000). Here, diversity is especially high in freshwater ecosystems, housing more than 6% of the global freshwater biodiversity, with particularly high numbers of endemic and rare taxa found on Mediterranean islands (De Figueroa et al. 2013). Yet, islands are also among the most threatened ecosystems within the region, with a predominance of non-perennial rivers and streams particularly vulnerable to anthropogenic impacts (Hopkins 2002; Skoulikidis et al. 2017).

Freshwater invertebrates, and in particular aquatic macroinvertebrates, are biological key components known to shape local communities and prime indicators of water quality. This also seems to be true in the streams on Mediterranean islands, where local macroinvertebrates are recognized as bioindicators of ecological disturbance (Erba et al. 2015; Feio et al. 2014; García et al. 2014). However, evidence about drivers influencing the observed insular aquatic macroinvertebrate diversity over time is scarce and restricted to a limited area (Garcia et al. 2017; Lobera et al. 2019). Studying their local spatio-temporal patterns of distribution is crucial for understanding the state and dynamics of freshwater ecosystems (Skoulikidis et al. 2017). Given that species diversity is high but taxonomic expertise, in particular also determination keys, are often not available, biological surveys and bioindication often rely on data from higher taxonomic levels only, typically family level. While this level increases reliability of the data, it also misses out the true diversity. DNA metabarcoding provides new opportunities to study diversity of freshwater communities (Carew et al. 2013; Elbrecht et al. 2017). In particular, DNA metabarcoding of environmental DNA (eDNA) provides a great tool for studying aquatic biodiversity in a non-invasive and time-efficient way (Deiner et al. 2016; Goldberg et al. 2015; Rees et al. 2014). While most aquatic eDNA surveys have focused on vertebrate species (Closek et al. 2019; Hänfling et al. 2016; Harper et al. 2018), recent studies have shown that a lot of information can also be obtained for invertebrate species (Leese et al. 2021; Macher et al. 2018; Mächler et al. 2019), despite the problem that the template molecules from the target taxa could be ‘watered down’ (Hajibabaei et al. 2019). Getting a broad picture of local macroinvertebrate communities from multiple fine-scale localities and from different time points within a single river system without investing a large amount of time in taxonomic identification could be of paramount importance for a better understanding of the observed diversity patterns as well as for its management (Stubbington et al. 2018).

Knowledge about the diversity and distribution of freshwater macroinvertebrates in the Mediterranean region is constantly growing, both due to multinational (e.g. Fauna Europaea, GBIF) and local initiatives (Ruffo and Stoch 2006), but is still missing for many regions. While the accessibility of reference DNA barcodes for macroinvertebrates is growing, there is still a significant gap present (Curry et al. 2018; Weigand et al. 2019). This holds true in particular for Mediterranean countries for which operational routine monitoring taxa lists often use higher taxonomic levels than species. Thus, given the incompleteness of the reference databases, taxonomic assignment of obtained DNA information can be troublesome. In taxonomic groups with even scarcer reference DNA sequences available like diatoms and other unicellular eukaryotes or meiofauna, where the majority of obtained information cannot be assigned to species level, there is a notion of using molecular operational taxonomic units (OTUs) (Feio et al. 2020; Westcott and Schloss 2015) or exact sequence variants (ESVs) (Tapolczai et al. 2019; Tapolczai et al. 2021) for analyses of diversity patterns. By often providing more ecologically relevant information (Tapolczai et al. 2021; Zizka et al. 2020) and higher reusability compared to OTUs (Schmidt et al. 2015), ESV-based approaches could provide a promising solution for studying remote and understudied regions, where much of the observed molecular diversity cannot be assigned to species level, yet.

In this study, we conducted the first eDNA survey for a stream ecosystem on Sicily. Sicily is the largest Mediterranean island and is located in the central part of Mediterranean basin. The freshwater macroinvertebrate fauna is relatively well documented with approximately 1300 species being reported from the insular freshwater ecosystems (Ruffo and Stoch 2006). The freshwater ecosystems on the island exhibit a comparatively high level of local endemism (Stoch 2000), in some groups like Malacostraca even more than 50% of taxa are endemic (Hupało et al. 2021). However, still very little is known about Sicilian freshwater macroinvertebrate communities and their local spatio-temporal distribution and dynamics. By focussing in detail on a single river system in the central part of Sicily, we aimed to provide insights into i) the general biodiversity patterns of macroinvertebrate communities, ii) the turnover of local macroinvertebrate communities between autumn and winter, and iii) whether dispersal in those communities is hindered by a barrier in the riverscape. Additionally, our aim was to compare patterns inferred from a species-based approach to an ESV-based approach. With that, we aimed to evaluate if eDNA metabarcoding data can provide valuable information for assessment and monitoring of Mediterranean freshwater biodiversity, even when the DNA reference databases are incomplete.

## Materials & Methods

### Field sampling

Water samples were collected during two seasonal sampling campaigns conducted in September 2019 (autumn) and January 2020 (winter) in the Fosso del Tempio river system on Sicily. The studied river system belongs to the basin of Fiume dei Monaci, which covers approximately 590 km^2^ and belongs to the Fiume Simeto catchment. The studied river originates from the slopes of Monte Moliano e Montagna on the border of the territory of the Municipalities of Aidone and Piazza Armerina. The main stretch of Fosso del Tempio is approximately 30 km long and after the first main confluence, it changes its name to Fosso Pietrarossa. To our knowledge there is no information regarding the average discharge of Fosso del Tempio.

A total of 27 sites were visited, with 18 being sampled both seasons, partially due to intermittency of parts of the studied river system, resulting in two sites (A6, E9) being completely dry during autumn sampling (Fig. 1, Tab. S1). To more broadly assess regional diversity, nine additional sites from neighbouring catchments were sampled at least in one season (Fig. 1). For sampling, a sterile 1 l bottle was used. Depending on the size of the stream, the water was taken either from the water surface at a single spot (1 l), or from each side of the stream (2x 0,5 l). All sampling was done against the flow direction wearing single-use nitrile gloves. Subsequently, samples were filtered on-site using a vacuum pump (Druck-Vakuumpumpe Typ:00A7.400 12V DC, Industrievertretung Neubauer) and a nitrocellulose filter (0.45 µm pore size, VWR, 513-1454). At each filtration location, a field blank (left open on the equipment for 30 seconds) was included and further processed with the other samples. If weather conditions did not allow for on-site filtration, samples were stored in a cooling box (4 °C) and filtered later during the same day (maximum 5 hours after sampling). If the filter clogged during filtration, a second, or third filter was used (Tab. S1). After filtration, the filters were folded with sterile tweezers and transferred into 1.5 ml Eppendorf tubes filled with 96% technical ethanol. Filters were immediately put into transportable ice boxes (<0°C) and within a few hours after sampling stored at -20 °C until DNA extraction.

**Fig. 1.**
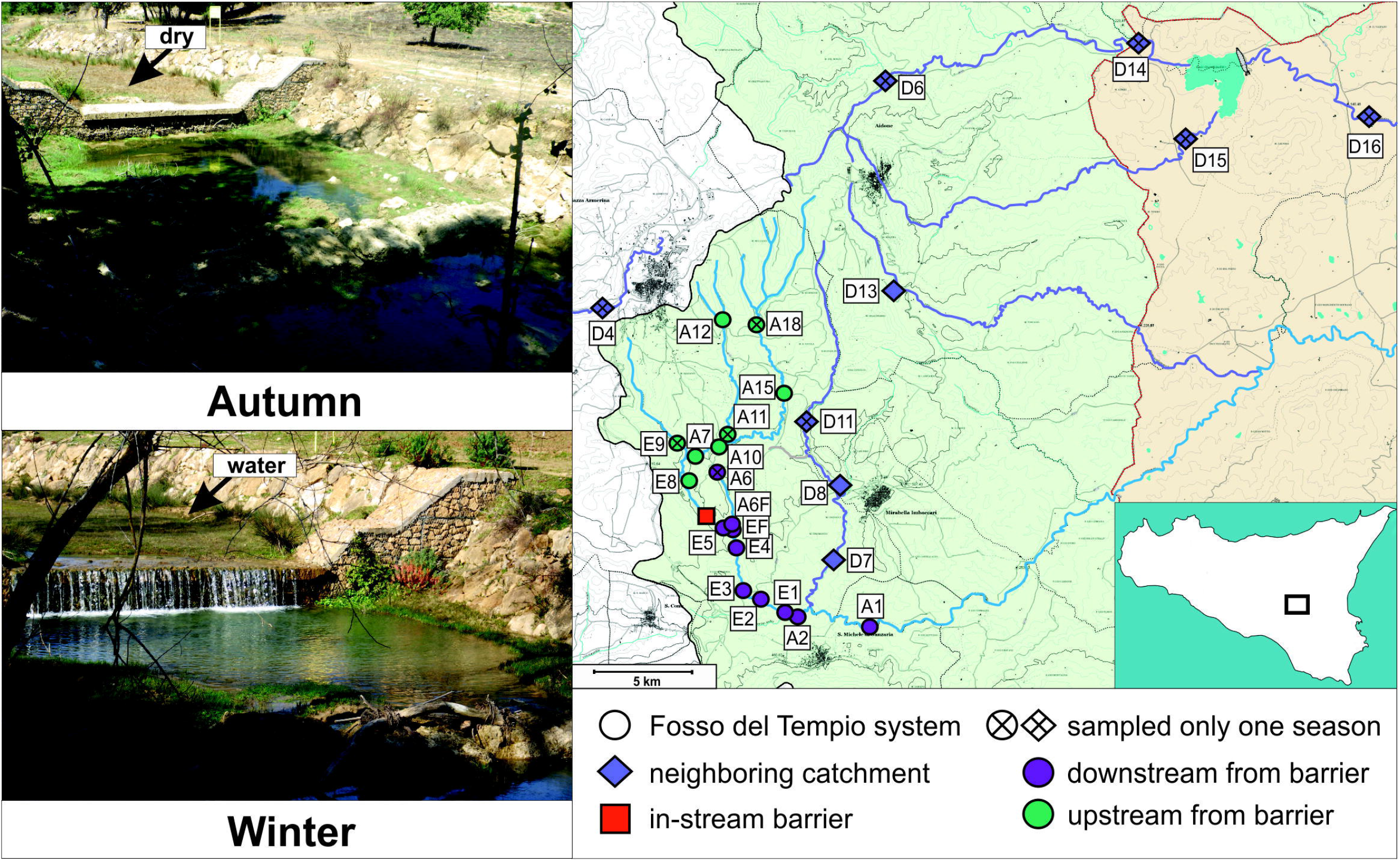
Map of sampling locations in the Fosso del Tempio system and neighbouring streams. Pictures on the left show sampling site E5 with the potential in-stream barrier in autumn and in winter.

### DNA extraction and purification

All laboratory work was conducted under sterile conditions in a specialized laboratory (‘eDNA lab’) that is only used for work on eDNA, regularly sterilized using UV light and with the air filtered with a HEPA 13 filter. Full body protective clothing was used (disposable overalls, facemasks, gloves, shoe covers) and the laboratory was irradiated with UV light prior to all working steps. Handling of the samples (DNA extraction; PCR) was conducted under independent UV hoods. First, the filters were taken from the ethanol and laid out in sterile petri dishes to dry for 15 to 20 hours. The petri dishes were covered with aluminium foil to protect them from UV irradiation in the eDNA lab. After drying, the filters were ripped into 6 to 7 pieces and transferred into 2 ml Eppendorf tubes containing 610 µl of lysis buffer (600 µl TNES and 10 µl Proteinase K (10 mg/ml)). DNA extraction was then conducted using a salt extraction protocol as described in Weiss and Leese (2016). Around 10 to 18 samples were extracted together. The field blanks were randomly distributed among sample batches and treated like normal samples in extractions.

Prior to DNA purification, 1 µl RNase (10 mg/ml, Thermoscientific, EN0531) was added to each sample and incubated at 37 °C for 30 minutes to remove RNA. After incubation, DNA was purified using the MinElute^®^ Reaction Cleanup Kit (Qiagen). All eDNA samples, where more than one filter had been used (Tab. S1) were pooled during purification by loading them onto the same column. After DNA purification, 68 DNA samples were retained for further processing (including negative filters). Purified DNA was resuspended in 20 µl nuclease-free H_2_O and stored at -20 °C. The DNA concentration was measured using the Qubit 2.0 (dsDNA High Sensitivity Array, Thermo Fisher Scientific, Beverly, USA)

### PCR amplification

A 205 bp long fragment of the cytochrome c oxidase subunit 1 (COI) mitochondrial gene was amplified using the fwh2 primer set (fwhF2 + fwhR2n) (Vamos et al. 2017) and the Multiplex PCR Plus Kit (100) (Qiagen GmbH). A two-step PCR protocol was employed, the first step with the untagged fwh2 primers and the second step with tagging primers (Leese et al. 2021).

To account for PCR stochasticity and to increase consistency of the results, two PCR replicates were processed for each sample (Zizka et al. 2019). In the first step, 1 µl of DNA was amplified in a 25 µl reaction (12.5 µl of the Multiplex Mastermix, 2.5 µl Color Dye, 0.25 µl fwh2F1 (10 mmol), 0.25 µl fwhR2n (10 mmol) and 8.5 µl PCR H_2_O). The cycling conditions for the first step were split into two parts, starting with 5 minutes at 95 °C for initial denaturation, followed by 10 cycles of a temperature step-down PCR (1°C per cycle) lasting 30 seconds going from 68 to 58°C, followed by final elongation for 30 seconds at 72 °C. The second part consisted of 25 cycles of 95°C for 30 sec, 58°C for 90 sec, and 72° for 30 sec. Final elongation was conducted for 10 minutes at 68 °C.

In the second PCR step, tagging primers were added. Each sample and each replicate were given a unique primer combination via a tag added to the PCR amplification primer to bioinformatically assign sequences to their original sample after sequencing (Tab. S2). The second PCR step was conducted with 1 µl of the the product from the first PCR step, 12.5 µl of the Multiplex Mastermix, 2.5 µl Coral Color Dye, 0.25 µl forward tagging primer (10 μM), 0.25 µl reverse tagging primer (10 μM) and 8.5 µl PCR H_2_O. Cycling conditions were as follows: 95°C for 5 minutes for initial denaturation, then 20 cycles of 95°C at 30 sec and 72°C for 120 sec with a final elongation at 68°C for 10 minutes.

The PCR success was checked using a 1% agarose gel and the products were measured with both the Qubit (2.0) High Sensitivity kit and the Fragment Analyzer™ Automated CE System (Advanced Analytical Technologies GmbH) using the NGS Standard Sensitivity Kit, before equimolar pooling (SequalPrep Normalization Plate, Applied Biosystems, Foster City, CA, USA). Residual primers were removed with a left sided SPRIselect size selection (Beckman Coulter, ratio: 0.76x). The final library was sent to Macrogen Europe and paired-end sequenced on one HiSeq X Illumina Lane (read length 2 × 150 bp). To increase sequence diversity, 5% PhiX were added by the sequencing company.

### Bioinformatic analyses

Raw reads for the two libraries were received from MacroGen as demultiplexed fastq files. The sequencing data was processed using a pre-release version of the graphical-user interface pipeline MetaProcessor (available at https://github.com/TillMacher/MetaProcessor). First, the quality of the raw reads was checked using FastQC (Andrews 2010). Subsequently, samples were renamed using a custom python script (Metaprocessor: rename_samples.py). Paired-end reads were merged using VSEARCH version 2.11.1 (Rognes et al. 2016), allowing for 25% differences between merged pairs and a minimum overlap of 20 bp. Afterwards, primers were trimmed using cutadapt version 2.8 (Martin 2011), using the linked adapter option without anchoring. Reads were then filtered by length (195-215 bp threshold for fwh2 target fragment) and by maximum expected error (maxee = 1), using VSEARCH. The filtered reads were dereplicated with singletons and chimeras removed with VSEARCH. All reads were then pooled using a custom python script and globally dereplicated (MetaProcessor: v_derep_singletons_uchime.py). Sequences were denoised into exact sequence variants (ESVs), using VSEARCH’s ‘--cluster_unoise’ function, with a minimum size of 8 and an alpha value of 2. The ESVs were remapped (usearch_global function, 100% similarity) to the individual sample files to create the read table. Taxonomic assignment of ESVs was conducted using BOLDigger version 1.1.10 (Buchner and Leese 2020) and the Barcode of Life data system (BOLD) database (Ratnasingham and Hebert 2007). The option “JAMP filter” was applied to extract the final taxonomy table.

Downstream processing of the dataset was conducted using TaxonTableTools (TTT) version 1.3.0 (Macher et al. 2021a). First, the full taxonomy table was processed for replicate consistency, where only ESVs present in both replicates were retained by using ‘merge replicates’ option. Afterwards, the resulting TaXon table was filtered using a read-based filter with 0.0005% threshold filtering. Using such a low threshold value was possible due to the high number of reads obtained combined with the dual indexing strategy (same index in i7 and i5 read) and lack of reads in negative controls, ensuring high reliability of data with preserving the records of rare taxa observed. The final filtering step included removal of taxonomically unmatched ESVs. General patterns of diversity were studied based on read proportion pie charts generated on phylum and order level. The resulting charts were superimposed on circular neighbor-joining phylogenetic trees based on p-distance generated with MEGA7 software (Kumar et al. 2016). To investigate seasonal patterns, the TaXon tables including autumn and winter samples were compared by generating Venn diagrams. In order to evaluate the effect of in-stream barrier, the TaXon tables were first converted to presence-absence (p/a) tables and subsequently, beta-diversity heatmaps and 3D Principal Coordinate Analysis (PCoA) charts were generated and the respective ANOSIM (analysis of similarities) R values were analysed. In principle, the ANOSIM is an analysis of variance using the average measures of dissimilarity (in this case Jaccard distances) in species composition, comparing them between and within given samples. Resulting test statistic value R usually range between 0 and 1, where values close to 1 generally support the sample dissimilarity contrary to values close to 0 which indicate no differences (Chapman and Underwood 1999; Clarke 1993). Finally, to visualize species distribution, Site Occupancy plots and Parallel Categories (ParCat) diagrams were produced.

### Datasets for analyses

The final dataset comprising all ESVs remaining after quality filtering (‘All ESVs’ dataset) was subsequently filtered for ESVs assigned to macroinvertebrates which are recognized as freshwater taxa, including also ones which are considered amphibiotic with a larval stage being confined to freshwater habitats (‘MZB ESVs’ dataset). Taxonomic filtering was performed according to taxa information stored in the Global Biodiversity Information Facility (GBIF) and World Register of Marine Species (WoRMS) databases. Freshwater macroinvertebrate taxa identified as potentially new for Sicily were cross-validated with the publicly available checklist of Italian Fauna (Latella et al. 2007) as well as species’ distribution maps in GBIF database (which, however, may also contain some unvalidated entries). Out of all ESVs assigned to freshwater macroinvertebrates, ESVs belonging to the same species were grouped together and represented by a single entry on species level for the final dataset (‘MZB species’ dataset). Furthermore, depending on the research question, the number of sampling sites for certain analyses varied. Thus, for analyses of the seasonal differences, only the sites, which were sampled both seasons were used (= 18 sites; Fig. 1). For analyses of possible barrier effects, only the sites from the Fosso del Tempio systems were used (highlighted in green and violet in Fig. 1), regardless if the site was visited both seasons or not, since the data for the barrier effect were analysed separately for each season. The downstream dataset consisted of nine sites of the Fosso del Tempio system, which were sampled in both seasons. The upstream dataset varied between seasons with seven sites being included in autumn and six in winter. There were five core upstream sites included in both seasons with three sites being sampled only in one season (A11, A18 in autumn and E9 in winter; Fig. 1). The downstream site A6, sampled only in winter, was excluded from the dataset given very low observed ESV/species numbers, pointing to potential strong effects of non-perennial flow, which could introduce a strong bias in barrier analysis. All filtering steps described above were conducted using taxon-based and sample-based filtering options in TTT software.

## Results

We obtained 445,683,202 raw read pairs, which also included reads of a second library with a different primer that was run in parallel on the same sequencing run. After paired-end merging, 232,498,448 reads were retrieved from fwhF2+fwhR2n primer pair. After the final quality filtering step 120,167,492 raw reads remained (31,900 reads assigned to negative controls), which were denoised into 7888 ESVs out of which 7745 could be taxonomically assigned. The subsequent merging of the PCR replicates retained 110,605,585 reads (no reads present in negative controls). The final read table (corresponding to ‘All ESVs’ dataset; Tab. S3) obtained after 0.0005% threshold filtering and removal of taxonomically unmatched ESVs, consisted of 110,148,185 reads, which were assigned to 7331 ESVs. Only around 6.5% (= 474) of all ESVs could be assigned to species level, resulting in 340 unique species. Taxonomic filtering for freshwater macroinvertebrates resulted in a read table consisting of 14,868,364 reads, assigned to 466 ESVs (‘MZB ESVs’ dataset; Tab. S4). Nearly 36% of macroinvertebrate ESVs (= 167) could be assigned to species level with 98 unique freshwater macroinvertebrate species retrieved.

### General patterns of diversity

The ESVs belonging to the ‘All ESVs’ dataset could be assigned to 29 phyla, with the highest proportion of reads as well as the highest number of ESVs being assigned to Arthropoda (Fig. 2). Apart from Arthropoda, high numbers of ESVs were retrieved for Heterokontophyta, Ochrophyta and Bacillariophyta. Altogether, above 40% of reads were assigned to non-metazoan taxa (Fig. 2). The majority of all species were assigned to Arthropoda, with Heterokontophyta and Annelida being the only other phyla with more than 10 species retrieved. Less than 0.5% of all reads consisted of 134 ESVs belonging to 15 phyla, both metazoan and non-metazoan.

**Fig. 2.**
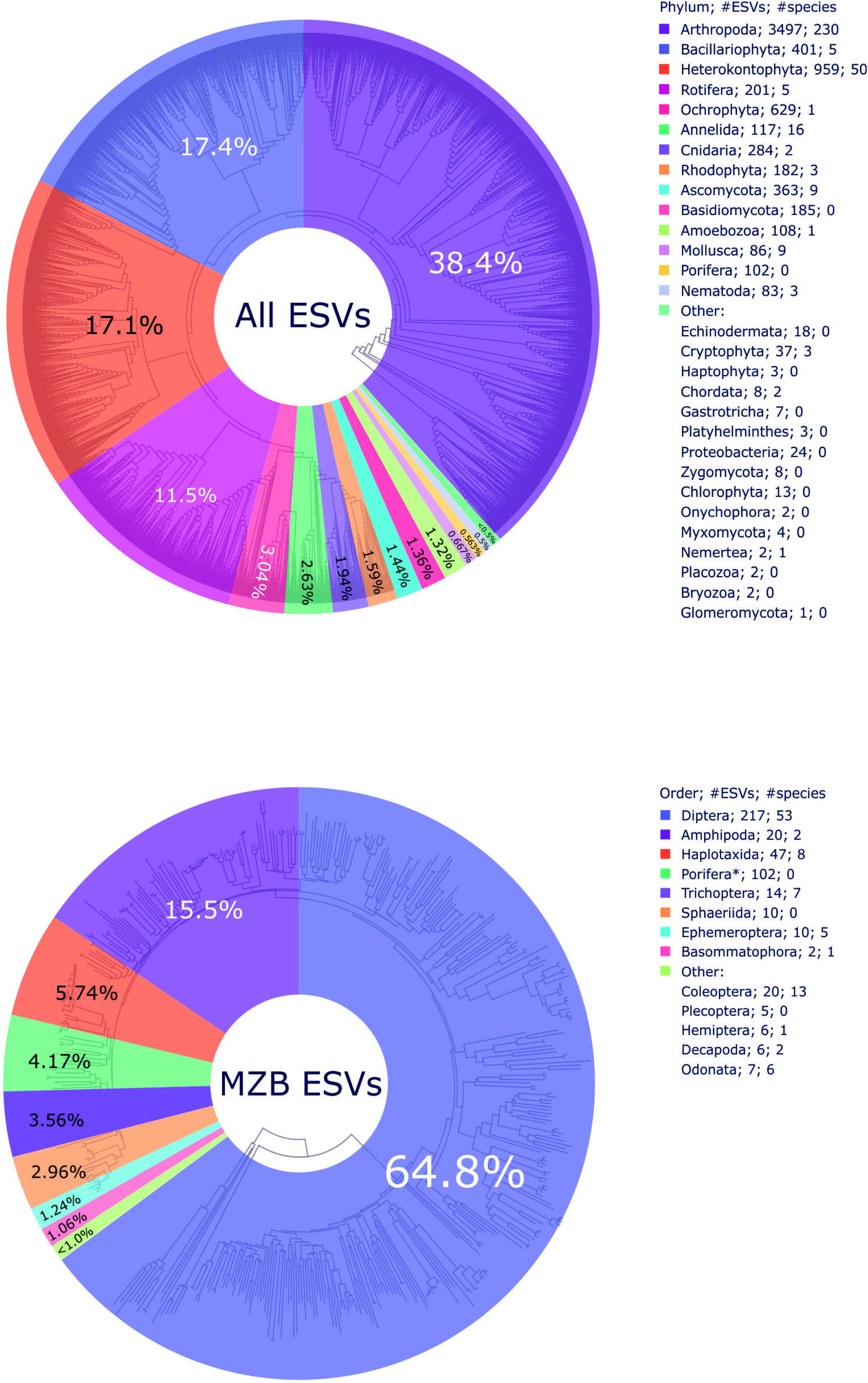
Read proportion charts with taxonomic composition, the number of ESVs assigned to a certain taxa group (phylum level for all ESVs; order level for freshwater macroinvertebrates*) and the number of unique species assigned to ESVs. Upper graph shows the read proportions for the entire dataset and lower graph read proportions in the freshwater macroinvertebrate dataset. *Since most ESVs assigned to Porifera could not be assigned to order level, they were kept to phylum level respectively.

The ESVs of the ‘MZB ESVs’ dataset were assigned to 13 orders, with nearly 65% of reads belonging to dipteran ESVs, which were also characterized by highest ESV and species diversity (Fig. 2). Next most read-abundant macroinvertebrate orders were amphipods and haplotaxidan annelids. More than 4% of reads were assigned to 102 poriferan ESVs, none of which could be assigned to species level with four being assigned to orders Bubarida and Desmacellida, respectively (Tab. S4). Nearly 10% of all macroinvertebrate ESVs (22 species of five orders) comprised less than 1% of reads.

#### Temporal turnover

The number of ESVs or species detected was higher in autumn than in winter regardless of the dataset, with a minor fraction of shared species/ESVs (Fig. 3). In the ‘All ESVs’ dataset, 4698 ESVs were retrieved in autumn and 2554 in winter, with 1288 ESVs being shared between seasons. In the ‘MZB ESVs’ dataset, 337 ESVs and 170 ESVs were retrieved in autumn and winter, respectively, with 118 shared between seasons. In the ‘MZB species’ dataset, 72 species were detected in autumn and 37 in winter, with 29 species found in both seasons. Less than half of the ESVs and macroinvertebrate species were found in winter compared to autumn, regardless of the dataset composition. The proportion of shared entities varied between approaches. In general, the percentage of shared diversity was inversely proportional to the broadness of the dataset, regardless of season analysed. In autumn, the proportion of diversity shared varied between roughly 27.4% for ‘All ESVs’ to 40.3% for ‘MZB species’, whereas in winter it ranged from 50.4% for ‘All ESVs’ to 78.4% for ‘MZB species’.

**Fig. 3.**
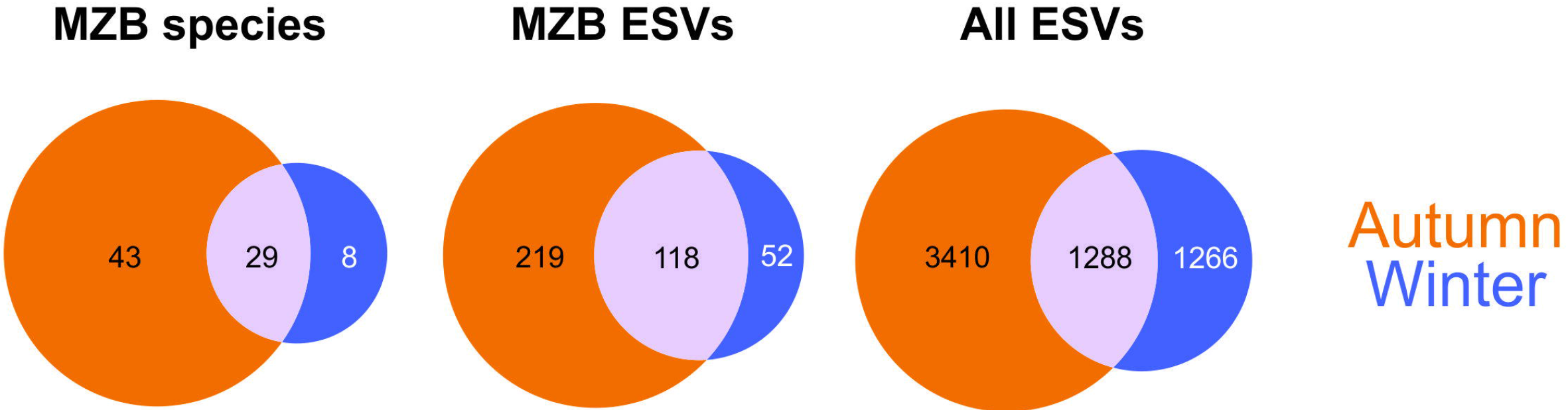
Venn diagrams showing number of species or ESVs detected in the autumn and winter samples as well as the fraction of shared species/ESVs between seasons for the three datasets analysed.

#### Barrier effect

We found strong community structuring across the barrier in autumn for all three datasets analysed (Fig. 4). ANOSIM R values (measure of average dissimilarity) were all highly significant (p<0.001) and ranged from 0.427 in ‘All ESVs’ dataset to 0.593 in ‘MZB ESVs’ dataset. In contrast, no significant differences were found for any of the datasets in winter, where ANOSIM R values varied from 0.08824 in ‘MZB species’ dataset up to 0.1565 in ‘MZB ESVs’ dataset (p < 0.05). This pattern was also reflected by the average Jaccard distances obtained per site (Fig. 4A, Tab. S5) as they were generally higher between sites across the barrier than on each side of the barrier regardless of season, except for upstream sites during winter. Considering individual sites, the Jaccard distances were on average higher across the barrier than within a stream section, with exception of two sites in autumn (A6F in ‘MZB ESVs’ and A1 in ‘All ESVs’ datasets) and 8 sites in winter (E3, E4, E5, A6F, E8, E9, A7 and A15 across various datasets; Tab. S5). However, the differences in the Jaccard distances between stream sections compared to sites within a stream section were more pronounced in autumn compared to winter. These results are further supported by the PCoA analyses performed on all sites within a stream section. In autumn clear separation between samples from downstream and upstream can be observed, when in winter there was no clear distinction between samples from the opposite sides of the in-stream barrier (Fig. 4B). Moreover, the direct comparison of sites closest to the in-stream barrier (E5 from the downstream and E8 from the upstream) indicate the proportion of diversity shared between those sites varies between seasons. In autumn, Jaccard distances ranged from 0.89 to 0.91, whereas in winter the values were between 0.67 and 0.76. The value of 1 obtained for those sites when using ‘MZB species’ dataset in winter should be treated with caution, since it is a result of no ESVs assigned to species level on site E5. A similar pattern can be observed in the proportion of shared diversity which ranges from 9% in case of ‘MZB ESVs’ to 10.5% for ‘MZB species’ whereas in winter it ranges from 23.5% in case of ‘MZB ESVs’ to 32.6% in ‘All ESVs’, with the exception of MZB species where none of seven ESV could be assigned to species level in site E5 (Fig. S1).

**Fig. 4.**
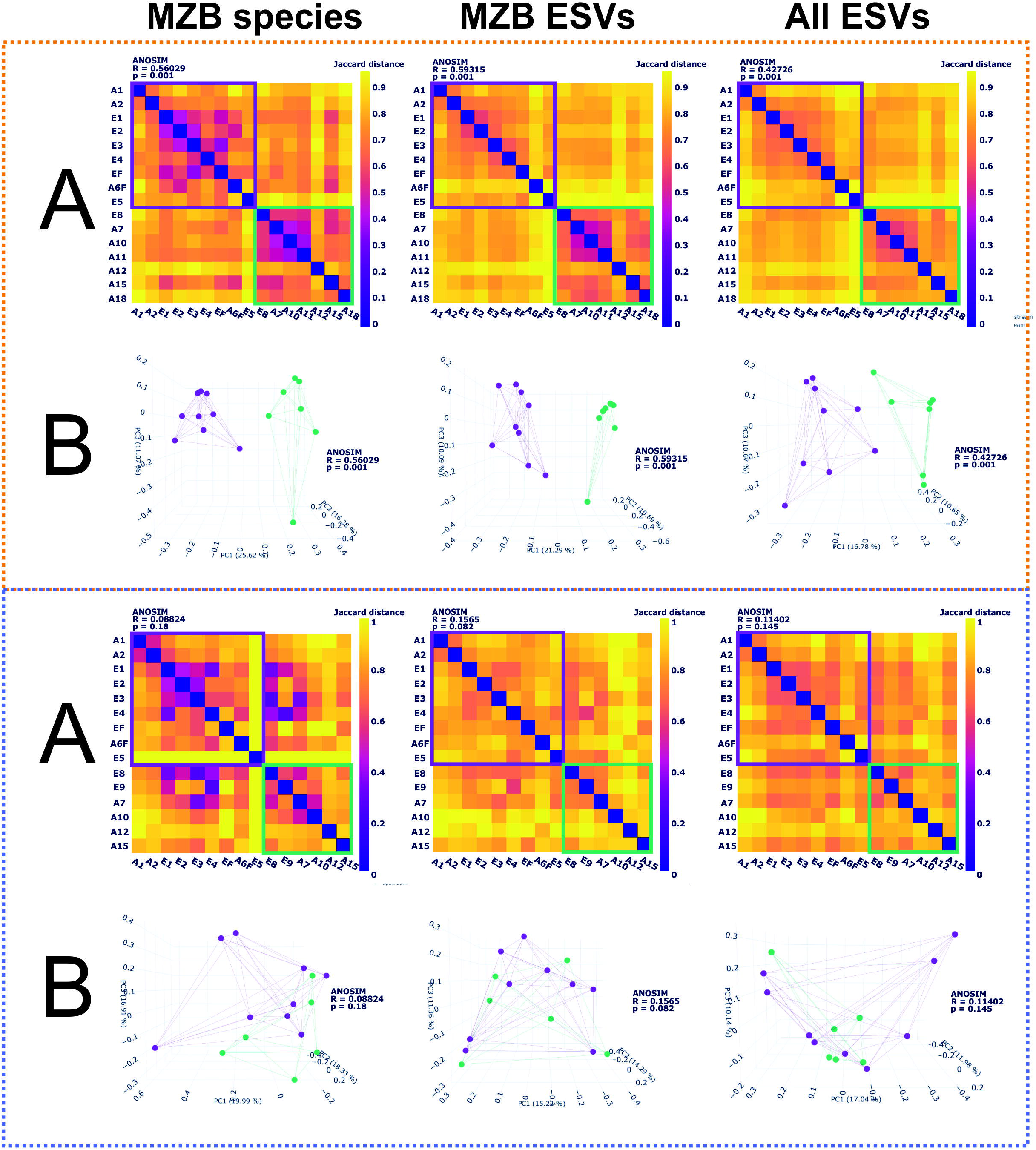
Diversity analyses evaluating the impact of the potential in-stream barrier in different seasons. A) Beta-diversity heatmaps. Bold squares indicate the stream section to which the samples belong (upstream/downstream of the barrier) B) Principal coordinate analysis (PCoA) graphs - full 3D versions available as html files in the Supplementary Material. The upper graphs (orange box) present the results for autumn, the lower graphs (blue box) present the results for winter. The color code corresponds to the one presented in Figure 1: violet = downstream sites, green = upstream sites.

### Macroinvertebrate species composition and distribution

Apart from looking at the diversity statistics, we also analysed the observed macroinvertebrate species composition as well as the seasonal turnover, including the potential effect of the barrier on species diversity. The distribution of the 98 freshwater macroinvertebrate species reported in the ‘MZB species’ dataset differed between seasons (Fig. 5A) and stream sections (Fig. 5B, C). Overall 32 of the 98 freshwater species were shared between Fosso del Tempio and neighboring streams, 52 species were exclusive to Fosso del Tempio and 14 to the neighbouring streams (Fig. S2). We identified 28 species potentially new for Sicily: 23 dipteran, 3 annelid, 1 caddisfly and 1 water beetle species, respectively (Tab. S6). Ten of them were found exclusively in the neighboring catchments.

**Fig. 5.**
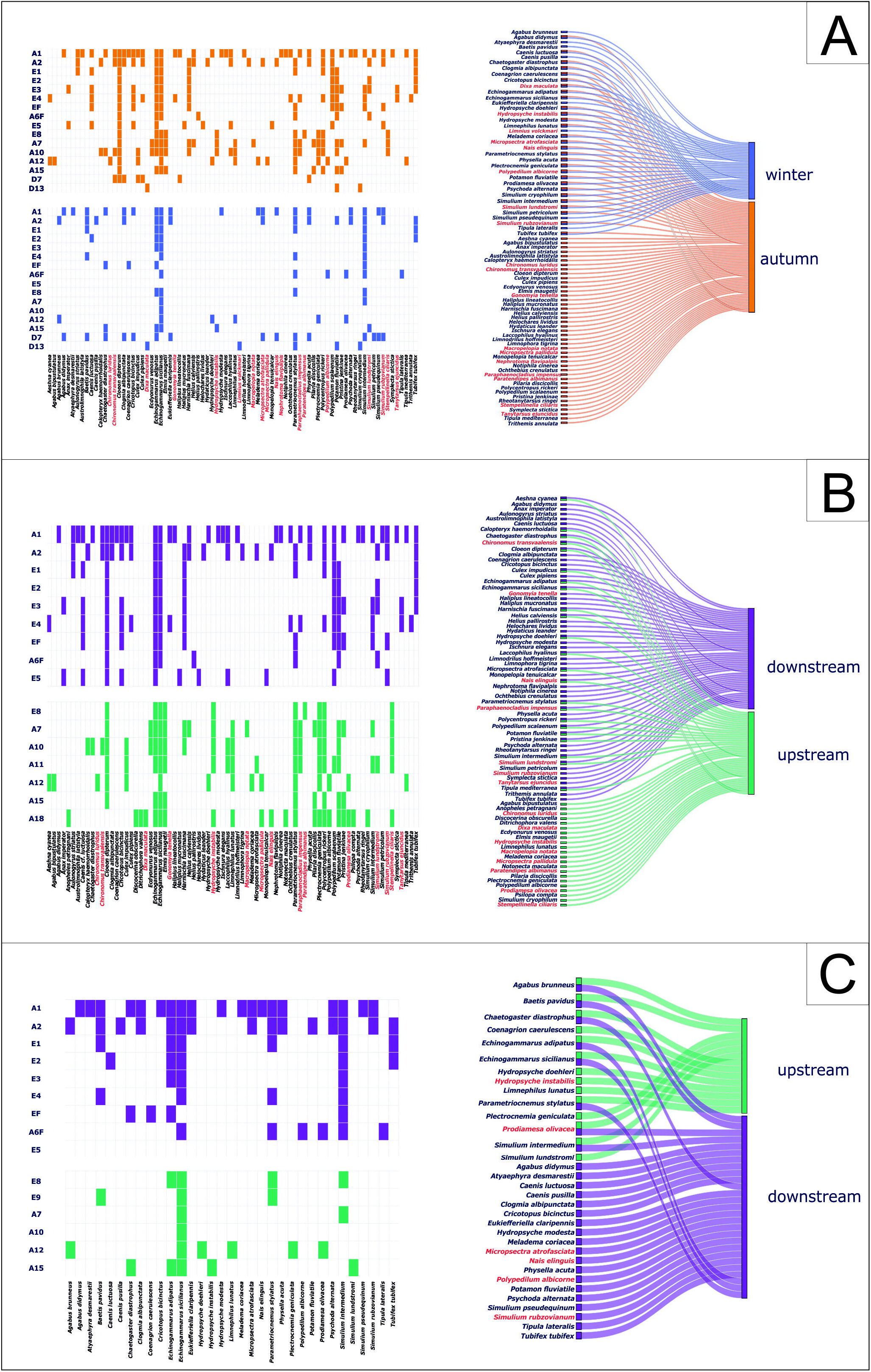
Macroinvertebrate species’ site occupancy plots (left) and ParCat distribution plots (right) for the ‘MZB species’ dataset. A) Species found in the two sampling seasons, B) Species sorted after stream section divided by the barrier - autumn, C) Species sorted after stream section divided by the barrier - winter. The species’ names marked in red indicate taxa potentially new for Sicily.

#### Temporal turnover

Regarding seasonality (Fig. 5A), there were 72 species found in autumn and 37 species found in winter with 29 retrieved in both seasons. Nearly half of all species found were dipterans. Over 40% of all species found in each season were only retrieved from a single site with mayfly *Cleon dipterum* being the most widespread species in autumn and *Echinogammarus sicilianus* being retrieved from most sites in winter. Notably both amphipod species and the only snail species were found both seasons along with nearly all caddisfly species (Tab. S7). On the contrary, most dragonflies and water beetles were observed in autumn, whereas two mayflies and a freshwater shrimp were retrieved only in winter.

#### Barrier effect

In autumn, 55 of the total 77 species were found in the downstream and 45 in the upstream section relative to the barrier, with 23 being shared between the sections (Fig. 5B). In the downstream sites eDNA detections of two species were retrieved from all sites: *Cleon dipterum* and *Echinogammarus adipatus*, whereas in the upstream *Echinogammarus sicilianus* and the chironomid *Parametriocnemus stylatus* were recorded from each site. Over 50% of the species detections were from single sites only. In winter, 28 of the total 33 species detected via eDNA in the Fosso del Tempio river system were found downstream of the barrier and 13 upstream with 8 shared between sections (Fig. 5C). Similarly to autumn, *Echinogammarus sicilianus* was retrieved from all sites in the upstream, whereas no species was recorded from all downstream sites with *Simulium intermedium* recorded from all sites but one. Again, in both stream sections the majority of species was recorded from single sites only. Interestingly, regardless of the season, most caddisfly species, along with only hemipteran, were found in the upstream sections only, whereas most dragonflies and water beetles were retrieved downstream of the barrier (Tab. S7). Notably both amphipod species were found on both sides of the barrier in both seasons.

## Discussion

### Observed diversity - freshwater macroinvertebrates and beyond

By using eDNA metabarcoding, we detected the presence of 98 freshwater macroinvertebrate species from the Fosso del Tempio river system and its neighboring catchment. The detected diversity counts for approximately 5% of all freshwater macroinvertebrate species known from the island (Latella et al. 2007; Ruffo and Stoch 2006). To the best of our knowledge, no freshwater macroinvertebrate biomonitoring data for Fosso del Tempio exist and thus, it is hard to estimate what fraction of the local diversity we were able to retrieve. However, given that a relatively small portion of Sicily’s hydrological network was covered with this study, the number of macroinvertebrate species retrieved is quite high. Moreover, taking into consideration that nearly 65% of obtained diversity could not be assigned to species level, the detected diversity is even higher. The majority of all freshwater macroinvertebrate species detected are arthropods, which were also the most dominant group in terms of read abundance. Among arthropods, the most dominant order in terms of both read abundance and diversity was Diptera. This result was to be expected since nearly half of all freshwater macroinvertebrates reported from Sicily belong to this order (Latella et al. 2007; Ruffo and Stoch 2006). It is also in agreement with other eDNA-based bioassessment of freshwater macroinvertebrates, where dipteran representatives are among the most dominant groups (Fernández et al. 2019; Leese et al. 2021). Within Diptera, we have detected 21 species belonging to the ecologically important family Chironomidae, which along with 12 species of ‘so-called’ EPT (Ephemeroptera, Plecoptera, Trichoptera) taxa, could serve as a basis for estimating the ecological state of the freshwater ecosystem in Fosso del Tempio river system (Wallace et al. 1996). Notably, we have also detected the presence of a single freshwater snail, the highly invasive snail *Physella (Physa) acuta*, known from Sicily already since the mid-19th century (Vinarski 2017). Since it was also the only snail species found, we have considered a possibility of primer bias towards inadequate amplification of molluscan DNA. However, after evaluating the primer binding for ten out of approximately 23 freshwater snail species known from Sicily (Ruffo and Stoch 2006), for which DNA information was available, we excluded that possibility since all of them could be hypothetically amplified with used primers (unpublished data). Absence of other snail taxa could be also resulting from low amounts of DNA shed in the environment either due to hard shell structure also possibly combined with very low numbers of individuals, which largely influence the detectability of an eDNA signal (Harrison et al. 2019; Stewart 2019). However, given we were still able to detect considerable diversity of taxa that bear hard carapax like crustaceans or water beetles, it could be rather due to the low number of individuals present. On the other hand, the observed low gastropod diversity could possibly indicate that invasive *P. acuta* have outcompeted other native snail species as already documented in other parts of the world (Dobson 2004; Zukowski and Walker 2009). This hypothesis is difficult to verify though, since no data on historical freshwater snail diversity in Fosso del Tempio is available. However, since *P. acuta* was only detected in relatively few samples, it might indicate either low abundance of snail species or low amount of DNA shed to the water.

In some cases, we also noticed a high level of intraspecific diversity, particularly in three species where more than 10 ESVs were ascribed to a single taxon namely a chironomid *Cricotopus bicinctus*, a simulid *Simulium rubzovianum* and an amphipod *Echinogammarus sicilianus*. Those findings confirm prior observation in all three taxa indicating already high degree of intraspecific diversity, in case of *E. sicilianus* the presence of potential cryptic diversity (Hupało et al. 2021; Sari et al. 2012; Sinclair and Gresens 2008). Interestingly, we have detected 28 species that to our knowledge, were not reported from Sicily before, indicating they might represent taxa potentially new for Sicily. Most of them belong to dipteran families Chironomidae, Simuliidae, Culicidae, Limoniidae, Dixidae, Empididae, Tipulidae and Muscidae with all species known to have an aquatic larval stage. Most of those species are known to occur in the Mediterranean basin indicating they might represent formerly overlooked valid records. However, five of them: *Culex inconspicuosus, Gonomyia tenella, Limnophora olympiae, Simulium ruficorne* and *Stempellinella ciliaris* were not reported from vicinity of Mediterranean basin with known occurrence data coming from Central and Northern Europe as well as Central and Southern Africa, indicating that those might represent false positive signals. Those false positives might result from mistakes present in public databases including incorrect synonyms and wrong sequence entries among others (Pentinsaari et al. 2020), which could contribute to observed records despite BOLDigger strategy to use the most common species across the most frequent of the most similar identification hits to enhance reliability of the record (Buchner and Leese 2020). Regarding other taxa, which might be new records for Sicily, there are three aquatic oligochaetes (*Dero furcata, Nais christinae, Nais elinguis*), all reported from the Mediterranean basin, one elmid beetle *Limnius volckmari* also occurring in the Mediterranean with a population on Corsica and one caddisfly *Hydropsyche instabilis* reported also from southern Italy. Although all of those occurrences might represent valid new records, one should treat them with extreme caution, since there were no actual specimens observed or collected from the sites, indicating that these require further investigation.

Our target organism group - freshwater macroinvertebrates - comprised 467 ESVs, which reflected about 6.5% of all ESV diversity obtained in this study. Although the vast majority of the remaining 6864 ESVs could not be assigned to species level, they still provide valuable information about the ecosystem surrounding the Fosso del Tempio system. We have retrieved traces of multiple entities of diatoms, other unicellular algae as well as heterotrophic and mixotrophic protists and fungi. Although only some could be assigned to species level, most of them could be resolved to higher taxonomic levels only, like phyla or classes. Since COI is suggested as barcoding region primarily for metazoans, in many cases the taxonomic assignment could probably be resolved better if other, group-specific molecular markers would be used e.g. rbcl, 18S rDNA or ITS (Evans et al. 2007; Pawlowski et al. 2012; Schoch et al. 2012). We have also retrieved DNA traces of several terrestrial arthropod species including 27 dipterans, 26 butterflies, 21 true bugs, 20 beetles, 12 hymenopterans, seven spiders, five orthopterans, five collembolans and a single mantid, phasmid, lacewing, bark lice and isopod. There was also a significant amount of information deriving from the freshwater organisms, which we did not consider as our target group. We have also observed DNA signals coming from the planktonic biota including five rotifers and two *Hydra* species. Moreover, the DNA trace of common carp (*Cyprinus carpio*) was retrieved from a single site. Interestingly, *C. carpio* established stable wild populations in inland waters of Sicily and is considered parautochthonous since being introduced supposedly by Romans (Marrone and Naselli-Flores 2015). It is likely representing a valid record since results obtained using fish-specific 12S marker (Taberlet et al. 2018) from the same sites confirmed its presence (Hupało et al. unpublished results). Altogether, the information included in the non-target part of the dataset obtained might provide valuable information about the surrounding ecosystem. More and more studies indicate the significance and validity of information contained in so-called bycatch derived from aquatic eDNA data (Macher et al. 2021b; Mariani et al. 2021). Thus, including and sharing the entire eDNA sequence dataset could be of paramount importance for further studies from the region and effort should be made towards their storage and public availability (Berry et al. 2020; Makiola et al. 2020). On the other hand, we have also retrieved a number of ESVs that represent obvious false positive records e.g. including ones that were taxonomically assigned to Echinodermata, a phylum that is known only to occur in marine waters. However, given the low level of genetic identity going below 80%, one could likely treat it as misidentification rather than a contamination, especially considering that no DNA signals were retrieved from negative controls. Regardless, one still should treat the records mentioned above with caution, because even though many of them could represent valid records, some of them might be deriving from sequencing errors and/or be a result of misidentification due to faultiness and incompleteness of reference databases.

### Insight into seasonality and barrier effect

Our results obtained with eDNA-based information provide an insight into the dynamics of the Fosso del Tempio river system. By conducting sampling campaigns in both autumn and winter seasons, we were able to investigate seasonal patterns of freshwater macroinvertebrate diversity. We observed more than twice as many macroinvertebrate species in autumn compared to winter. This result is to some degree unexpected. Even though differences in species composition in the Mediterranean region could be partially attributed to a high degree of water level fluctuation observed between arid summer and wet winter, the species richness is expected to remain similar between seasons (Bonada and Resh 2013; Gasith and Resh 1999). Given that samples were collected in September, at the end of dry season in Sicily and in January, which marks approximately the middle of wet season, the habitat and resource availability for aquatic biota are very different. This should reflect the changes in species’ assemblages and dominances, however the majority of species should be evolutionarily adapted to those varying conditions, resulting in certain similarities in species diversity between seasons (Gasith and Resh 1999). One of the reasons behind observed patterns could be visibly higher water levels during winter sampling. This could also have an indirect effect on lower diversity observed with the DNA concentration being more diluted and thus, less taxa being retrieved. That being said, with a relatively high number of species shared between seasons, one may assume that at least some of taxa reported only from autumn could in fact also occur on sites in winter and could be retrieved if a higher water volume was sampled accordingly. On the other hand, there were eight species that were only retrieved from winter samples. Although in some cases like *Eukieferella claripennis, Limnius volckmari* and *Tipula lateralis*, it seems to be the case of rarity of those taxa in autumn since they were only retrieved locally, other cases might show some true seasonal patterns. In case of mayflies *Baetis pavidus* and *Caenis pusilla* a possible niche overlap with *Caenis luctuosa* mediated with supposed seasonal fluctuations might be the reason behind observed species presence in close-by sites in autumn and winter. The same could be true for the water beetle *Agabus brunneus* ‘replacing’ *Agabus bipustulatus* on site A12 in winter compared to autumn. Interestingly, we have also retrieved an eDNA signal from freshwater shrimp *Atyaephyra desmarestii*, native to Mediterranean region. The retrieved information from winter eDNA sampling corresponds with the observations made in the field where numerous specimens were observed only during winter sampling with no specimen collected in autumn. Although the species is known to have strong seasonal fluctuations (Dhaouadi-Hassen and Boumaiza 2009; Fidalgo et al. 2015; Schoolmann et al. 2015), the species’ abundance should be higher in autumn compared to winter. Since the opposite is the case here, this puzzling observation likely requires further investigation.

Our data also indicate that the concrete weir present in the stream course might have an effect on dispersal of resident aquatic taxa. There are numerous studies confirming that weirs may act as barriers for macroinvertebrate dispersal significantly reducing both connectivity and observed species diversity on the opposite sites of the barrier (Brooks et al. 2018; Poff and Zimmerman 2010). The effect seems to be even more profound when there is a high degree of water level fluctuations, which is often the case in Mediterranean streams. Here, we observed a significant change of water level and flow velocity between dry and wet seasons. During sampling conducted in September we have seen virtually no water going through the weir, whereas in winter with the higher water level the change in amount of water flowing through was clearly visible, although flow was still disrupted by the barrier. This effect seems to be reflected by the results of similarity-based analyses with a clear distinction between the observed diversity in the downstream and upstream samples during dry season compared to wet season, where no visible difference can be observed. This is also reflected in the proportion of species that were found only in the upstream catchment between seasons with a much lower percentage of species uniquely found in the upstream in winter compared to autumn. A similar pattern was found when comparing diversity observed on two border sites separated by the barrier, where a comparatively higher proportion of diversity was shared between those sites in winter compared to autumn. Although, any species-related comparisons between the seasons has to be taken with caution due to various reasons described above, the observed effect of an in-stream weir could have significant implications for observed diversity in the Fosso del Tempio system. Based on our results, the weir seems to hinder the connectivity between upstream and downstream parts of the system with a portion of taxa being found on both sides of the barrier. Regardless of season, the majority of taxa shared between stream sections are the hemilimnic ones with flying adult stages. Increased dispersal ability in flying insects with aquatic larvae has been proven to be important in terms of observed patterns of diversity in a stream ecosystem with high degree of similarity between river sections regardless of any possible barriers (Hughes et al. 2009). Similarly, the opposite is true for the species with lower dispersal abilities with more fractionated and restricted patterns of genetic diversity. In such cases, the in-stream barriers might lead to genetic differentiation leading to strong population structuring and in some cases, to allopatric speciation. We have observed a similar case in our data, where certain ESVs of *Echinogammarus sicilianus*, a freshwater amphipod with limited dispersal abilities, are present only in the downstream or upstream section of the Fosso del Tempio, regardless of the season. This finding might indicate that the intraspecific diversity observed might, at least partially, be resulting from the effect of the in-stream weir.

### Different approaches, similar results - ESV vs. species comparability

The results regarding the seasonality patterns and the effect of the in-stream barrier obtained with macroinvertebrate species-based approach were similar to the ones obtained with all macroinvertebrate ESVs including more than 60% of all macroinvertebrate ESVs which could not be assigned to species level. Although notable progress in DNA barcoding of aquatic biota has been made in recent years, there is still a significant amount of data missing in reference databases (Weigand et al. 2019). The difference in freshwater macroinvertebrate barcode coverage varies between different regions due to unbalanced efforts taken in filling the missing gaps. Our results indicate that Sicily freshwater macroinvertebrates seem to be relatively underrepresented with only 35% of our dataset being assigned to species level, compared to European average of approximately 64.5% freshwater macroinvertebrate species having at least a single barcode publicly available (Weigand et al. 2019). In those cases, ESV-based approaches could provide an alternative solution for studying freshwater ecosystem dynamics. This seems to be also of particular importance for research based on organisms where taxonomy is complex and not fully resolved like diatoms where the use of ESVs has been successfully implemented (Tapolczai et al. 2019). Moreover, considering ESVs provides a finer level of detail to observed diversity by providing an insight also into intraspecific diversity, which is discarded when species or clusters serving as species proxy (e.g. OTUs) are considered (Tapolczai et al. 2021; Zizka et al. 2020). This could be of paramount importance when considering phylogeography or population genetics of observed taxa (Antich et al. 2021; Turon et al. 2020).

Since the primers used in this study amplify a broad range of taxa groups and given that macroinvertebrate diversity represented only a fraction of all retrieved ESVs, we have also decided to look at the results obtained with an approach where all generated diversity will be taken into consideration. Surprisingly, despite taxonomic broadness of the dataset, including also a high portion of non-freshwater biota, the results obtained were similar to those obtained only with macroinvertebrate data. This finding seems particularly interesting pointing out the high sensitivity and reproducibility of generated results. Even though the results deriving from this general approach should be looked at with increased awareness, they still can inform about community connectivity or seasonality patterns in a stream ecosystem.

## Conclusions

Our results provide a first insight into freshwater diversity and community dynamics of the Fosso del Tempio river system in Sicily, used as an exemplary Mediterranean insular freshwater ecosystem. We showcase the potential that comes with using eDNA metabarcoding for studying freshwater ecosystems, even in understudied regions with still significant portions of genetic information missing like the Mediterranean islands. Based on our findings, an ESV-based approach provides a promising, highly reproducible approach for studying freshwater community patterns and dynamics even - or in particular - in highly understudied regions. By providing reproducible taxonomic units, data could be easily reused in the future with new regional biodiversity data added and with increasing completeness of local reference databases. Hence, we highlight the need for a unified digital storage and access solution, which could ensure the public availability of the data at both regional and international level. Specifically for regions with non-perennial flows, hydrological differences between seasons need to be considered and incorporated in bioassessment planning, sampling and interpretation. We stress the importance of proper sampling design taking into consideration seasonality and equal representation of sites across the entire river system to maximize the representation of the regional community and its dynamics.

## Declarations

### Funding

This research was funded by German Research Foundation (DFG) (project LE 2323/9-1).

### Conflicts of interest

None declared

### Availability of data and material

Data is publicly available in the European Nucleotide Archive (PRJEB45583).

### Code availability

MetaProcessor; available at https://github.com/TillMacher/MetaProcessor

### Ethics approval

Not applicable

### Consent to participate and publication

All authors agreed to participate in publication process.

## Data Accessibility

The data is publicly available in the European Nucleotide Archive (PRJEB45583).

## Acknowledgements

We would like to thank Marie-Thérése Werner and Pedro M.G. Gomes for their valuable help during the sampling campaigns. We would also like to thank Fabio Stoch for providing in-depth information about the studied river system. We also thank Bernd Sures, Daniel Grabner, Diego Fontaneto for the logistic support and Florian Altermatt, Michał Grabowski, Alexander Weigand and Hannah Weigand for helpful discussions throughout the project.

This research was funded by German Research Foundation (DFG) (project LE 2323/9-1).

## Figure captions

**Fig. S1** Venn diagrams showing the shared number of ESVs/species between the two sites closest to and divided by the in-stream barrier, according to the season. Violet = sampling site E5 (downstream from the barrier), green = sampling site E8 (upstream from the barrier).

**Fig. S2** ParCat distribution plot of macroinvertebrate species in the Fosso del Tempio river system and the neighboring streams.

**Tab. S1** The list of sampling localities. The names in bold indicate samples belonging to the Fosso del Tempio river system. Sites marked with an asterisk were sampled in a single season only.

**Tab. S2** i5 and i7 indices used for each sample and replicate in 2nd step PCR, and number of raw and filtered (replicates merged) reads obtained after sequencing.

**Tab. S3** Complete TaXon table obtained after final filtering steps (‘All ESVs’ dataset).

**Tab. S4** TaXon table obtained after taxonomic filtering for freshwater macroinvertebrates (‘MZB ESVs’ dataset and ‘MZB species’).

**Tab. S5** Jaccard distances for different seasons and different datasets. MeanW values indicate mean values for a particular site within the stream section the site belongs to, whereas MeanB values indicate mean values between stream sections. ∑ values indicate mean values obtained for all sites within particular stream section.

**Tab. S6** TaXon table with the macroinvertebrate species potentially new for Sicily.

**Tab. S7** Number of macroinvertebrate species per order, according to the seasonality and the in-stream barrier.

## Notes

### Competing Interest Statement

The authors have declared no competing interest.

## References

Almond, R., M. Grooten & T. Peterson, 2020. Living Planet Report 2020 - Bending the curve of biodiversity loss. World Wildlife Fund.

Andrews, S., 2010. FastQC: a quality control tool for high throughput sequence data. vol Available online at: https://www.bioinformatics.babraham.ac.uk/projects/fastqc/. Babraham Bioinformatics, Babraham Institute, Cambridge, United Kingdom.

Antich, A., C. Palacin, O. S. Wangensteen & X. Turon, 2021. To denoise or to cluster, that is not the question: optimizing pipelines for COI metabarcoding and metaphylogeography. BMC bioinformatics 22(1):1–24.

Arnell, N. W., 1999. Climate change and global water resources. Global environmental change 9:S31–S49.

Balian, E. V., H. Segers, K. Martens & C. Lévéque, 2007. The freshwater animal diversity assessment: an overview of the results Freshwater animal diversity assessment. Springer, 627–637.

Berry, O., S. Jarman, A. Bissett, M. Hope, C. Paeper, C. Bessey, M. K. Schwartz, J. Hale & M. Bunce, 2020. Making environmental DNA (eDNA) biodiversity records globally accessible. Environmental DNA.

Bonada, N. & V. H. Resh, 2013. Mediterranean-climate streams and rivers: geographically separated but ecologically comparable freshwater systems. Hydrobiologia 719(1):1–29.

Brooks, A. J., B. Wolfenden, B. J. Downes & J. Lancaster, 2018. Barriers to dispersal: The effect of a weir on stream insect drift. River Research and Applications 34(10):1244–1253.

Buchner, D. & F. Leese, 2020. BOLDigger–a Python package to identify and organise sequences with the Barcode of Life Data systems. Metabarcoding and Metagenomics 4:e53535.

Carew, M. E., V. J. Pettigrove, L. Metzeling & A. A. Hoffmann, 2013. Environmental monitoring using next generation sequencing: rapid identification of macroinvertebrate bioindicator species. Frontiers in zoology 10(1):1–15.

Chapman, M. & A. Underwood, 1999. Ecological patterns in multivariate assemblages: information and interpretation of negative values in ANOSIM tests. Marine ecology progress series 180:257–265.

Clarke, K. R., 1993. Non-parametric multivariate analyses of changes in community structure. Australian journal of ecology 18(1):117–143.

Closek, C. J., J. A. Santora, H. A. Starks, I. D. Schroeder, E. A. Andruszkiewicz, K. M. Sakuma, S. J. Bograd, E. L. Hazen, J. C. Field & A. B. Boehm, 2019. Marine vertebrate biodiversity and distribution within the central California Current using environmental DNA (eDNA) metabarcoding and ecosystem surveys. Frontiers in Marine Science 6:732.

Curry, C. J., J. F. Gibson, S. Shokralla, M. Hajibabaei & D. J. Baird, 2018. Identifying North American freshwater invertebrates using DNA barcodes: are existing COI sequence libraries fit for purpose? Freshwater Science 37(1):178–189.

De Figueroa, J. M. T., M. J. López-Rodríguez, S. Fenoglio, P. Sánchez-Castillo & R. Fochetti, 2013. Freshwater biodiversity in the rivers of the Mediterranean Basin. Hydrobiologia 719(1):137–186.

Deiner, K., E. A. Fronhofer, E. Mächler, J.-C. Walser & F. Altermatt, 2016. Environmental DNA reveals that rivers are conveyer belts of biodiversity information. Nature communications 7(1):1–9.

Dhaouadi-Hassen, S. & M. Boumaiza, 2009. Reproduction and population dynamics of Atyaephyra desmarestii (Decapoda, Caridea) from the Sidi Salem dam lake (northern Tunisia). Crustaceana:129–139.

Dobson, M., 2004. Replacement of native freshwater snails by the exotic Physa acuta (Gastropoda: Physidae) in southern Mozambique; a possible control mechanism for schistosomiasis. Annals of Tropical Medicine & Parasitology 98(5):543–548.

Dudgeon, D., A. H. Arthington, M. O. Gessner, Z.-I. Kawabata, D. J. Knowler, C. Lévêque, R. J. Naiman, A.-H. Prieur-Richard, D. Soto & M. L. Stiassny, 2006. Freshwater biodiversity: importance, threats, status and conservation challenges. Biological reviews 81(2):163–182.

Elbrecht, V., E. E. Vamos, K. Meissner, J. Aroviita & F. Leese, 2017. Assessing strengths and weaknesses of DNA metabarcoding-based macroinvertebrate identification for routine stream monitoring. Methods in Ecology and Evolution 8(10):1265–1275.

Erba, S., G. Pace, D. Demartini, D. Di Pasquale, G. Dörflinger & A. Buffagni, 2015. Land use at the reach scale as a major determinant for benthic invertebrate community in Mediterranean rivers of Cyprus. Ecological Indicators 48:477–491.

Evans, K. M., A. H. Wortley & D. G. Mann, 2007. An assessment of potential diatom “barcode” genes (cox1, rbcL, 18S and ITS rDNA) and their effectiveness in determining relationships in Sellaphora (Bacillariophyta). Protist 158(3):349–364.

Feio, M., J. Ferreira, A. Buffagni, S. Erba, G. Dörflinger, M. Ferréol, A. Munné, N. Prat, I. Tziortzis & G. Urbanič, 2014. Comparability of ecological quality boundaries in the Mediterranean basin using freshwater benthic invertebrates. Statistical options and implications. Science of the Total Environment 476:777–784.

Feio, M. J., S. R. Serra, A. Mortágua, A. Bouchez, F. Rimet, V. Vasselon & S. F. Almeida, 2020. A taxonomy-free approach based on machine learning to assess the quality of rivers with diatoms. Science of the Total Environment 722:137900.

Fernández, S., S. Rodríguez-Martínez, J. L. Martínez, E. Garcia-Vazquez & A. Ardura, 2019. How can eDNA contribute in riverine macroinvertebrate assessment? A metabarcoding approach in the Nalón River (Asturias, Northern Spain). Environmental DNA 1(4):385–401.

Fidalgo, M. L., P. Santos, C. Ferreira & A. Silva, 2015. Population structure and dynamics of the freshwater shrimp Atyaephyra desmarestii (Millet, 1831) in the lower River Minho (NW Portugal). Crustaceana 88(6):657–673.

Garcia, C., C. N. Gibbins, I. Pardo & R. J. Batalla, 2017. Long term flow change threatens invertebrate diversity in temporary streams: Evidence from an island. Science of the Total Environment 580:1453–1459.

García, L., I. Pardo & C. Delgado, 2014. Macroinvertebrate indicators of ecological status in Mediterranean temporary stream types of the Balearic Islands. Ecological indicators 45:650–663.

Gasith, A. & V. H. Resh, 1999. Streams in Mediterranean climate regions: abiotic influences and biotic responses to predictable seasonal events. Annual review of ecology and systematics 30(1):51–81.

Goldberg, C. S., K. M. Strickler & D. S. Pilliod, 2015. Moving environmental DNA methods from concept to practice for monitoring aquatic macroorganisms. Biological Conservation 183:1–3.

Grill, G., B. Lehner, M. Thieme, B. Geenen, D. Tickner, F. Antonelli, S. Babu, P. Borrelli, L. Cheng & H. Crochetiere, 2019. Mapping the world’s free-flowing rivers. Nature 569(7755):215–221.

Hajibabaei, M., T. M. Porter, C. V. Robinson, D. J. Baird, S. Shokralla & M. T. Wright, 2019. Watered-down biodiversity? A comparison of metabarcoding results from DNA extracted from matched water and bulk tissue biomonitoring samples. PloS one 14(12):e0225409.

Hänfling, B., L. Lawson Handley, D. S. Read, C. Hahn, J. Li, P. Nichols, R. C. Blackman, A. Oliver & I. J. Winfield, 2016. Environmental DNA metabarcoding of lake fish communities reflects long-term data from established survey methods. Molecular ecology 25(13):3101–3119.

Harper, L. R., L. Lawson Handley, C. Hahn, N. Boonham, H. C. Rees, K. C. Gough, E. Lewis, I. P. Adams, P. Brotherton & S. Phillips, 2018. Needle in a haystack? A comparison of eDNA metabarcoding and targeted qPCR for detection of the great crested newt (Triturus cristatus). Ecology and evolution 8(12):6330–6341.

Harrison, J. B., J. M. Sunday & S. M. Rogers, 2019. Predicting the fate of eDNA in the environment and implications for studying biodiversity. Proceedings of the Royal Society B 286(1915):20191409.

Hopkins, L., 2002. IUCN and the Mediterranean Islands: Opportunities for biodiversity conservation and sustainable use. International Union for Conservation of Nature: Gland, Switzerland.

Hughes, J. M., D. J. Schmidt & D. S. Finn, 2009. Genes in streams: using DNA to understand the movement of freshwater fauna and their riverine habitat. BioScience 59(7):573–583.

Hupało, K., F. Stoch, I. Karaouzas, A. Wysocka, T. Rewicz, T. Mamos & M. Grabowski, 2021. Freshwater Malacostraca of the Mediterranean Islands–Diversity, Origin, and Conservation Perspectives Recent Advances in Freshwater Crustacean Biodiversity and Conservation. CRC Press, 139–220.

Kumar, S., G. Stecher & K. Tamura, 2016. MEGA7: molecular evolutionary genetics analysis version 7.0 for bigger datasets. Molecular biology and evolution 33(7):1870–1874.

Latella, L., S. Ruffo & F. Stoch, 2007. The project CKmap (Checklist and distribution of the Italian fauna) methods and informatical techniques. Memorie del Museo Civico di Storia Naturale di Verona 17:15–19.

Leese, F., M. Sander, D. Buchner, V. Elbrecht, P. Haase & V. M. Zizka, 2021. Improved freshwater macroinvertebrate detection from environmental DNA through minimized nontarget amplification. Environmental DNA.

Lobera, G., I. Pardo, L. García & C. García, 2019. Disentangling spatio-temporal drivers influencing benthic communities in temporary streams. Aquatic Sciences 81(4):1–17.

Macher, J. N., A. Vivancos, J. J. Piggott, F. C. Centeno, C. D. Matthaei & F. Leese, 2018. Comparison of environmental DNA and bulk-sample metabarcoding using highly degenerate cytochrome c oxidase I primers. Molecular ecology resources 18(6):1456–1468.

Macher, T.-H., A. J. Beermann & F. Leese, 2021a. TaxonTableTools-A comprehensive, platform-independent graphical user interface software to explore and visualise DNA metabarcoding data. Molecular Ecology Resources.

Macher, T.-H., R. Schütz, J. Arle, A. J. Beermann, J. Koschorreck & F. Leese, 2021b. Beyond fish eDNA metabarcoding: Field replicates disproportionately improve the detection of stream associated vertebrate species. bioRxiv.

Mächler, E., C. J. Little, R. Wüthrich, R. Alther, E. A. Fronhofer, I. Gounand, E. Harvey, S. Hürlemann, J. C. Walser & F. Altermatt, 2019. Assessing different components of diversity across a river network using eDNA. Environmental DNA 1(3):290–301.

Makiola, A., Z. G. Compson, D. J. Baird, M. A. Barnes, S. P. Boerlijst, A. Bouchez, G. Brennan, A. Bush, E. Canard & T. Cordier, 2020. Key questions for next-generation biomonitoring. Frontiers in Environmental Science 7:197.

Mariani, S., L. R. Harper, R. A. Collins, C. Baillie, O. Wangensteen, A. McDevitt, M. Heddell-Cowie & M. J. Genner, 2021. Estuarine molecular bycatch as a landscape-wide biomonitoring tool. bioRxiv.

Marrone, F. & L. Naselli-Flores, 2015. A review on the animal xenodiversity in Sicilian inland waters (Italy). Advances in Oceanography and Limnology.

Martin, M., 2011. Cutadapt removes adapter sequences from high-throughput sequencing reads. EMBnet journal 17(1):10–12.

Médail, F. & P. Quézel, 1999. Biodiversity hotspots in the Mediterranean Basin: setting global conservation priorities. Conservation biology 13(6):1510–1513.

Myers, N., R. A. Mittermeier, C. G. Mittermeier, G. A. Da Fonseca & J. Kent, 2000. Biodiversity hotspots for conservation priorities. Nature 403(6772):853.

Pawlowski, J., S. Audic, S. Adl, D. Bass, L. Belbahri, C. Berney, S. S. Bowser, I. Cepicka, J. Decelle & M. Dunthorn, 2012. CBOL protist working group: barcoding eukaryotic richness beyond the animal, plant, and fungal kingdoms. PLoS Biol 10(11):e1001419.

Pentinsaari, M., S. Ratnasingham, S. E. Miller & P. D. Hebert, 2020. BOLD and GenBank revisited–Do identification errors arise in the lab or in the sequence libraries? PloS one 15(4):e0231814.

Poff, N. L. & J. K. Zimmerman, 2010. Ecological responses to altered flow regimes: a literature review to inform the science and management of environmental flows. Freshwater biology 55(1):194–205.

Ratnasingham, S. & P. D. Hebert, 2007. BOLD: The Barcode of Life Data System (http://www.barcodinglife.org). Molecular ecology notes 7(3):355–364.

Rees, H. C., B. C. Maddison, D. J. Middleditch, J. R. Patmore & K. C. Gough, 2014. The detection of aquatic animal species using environmental DNA–a review of eDNA as a survey tool in ecology. Journal of Applied Ecology 51(5):1450–1459.

Reid, A. J., A. K. Carlson, I. F. Creed, E. J. Eliason, P. A. Gell, P. T. Johnson, K. A. Kidd, T. J. MacCormack, J. D. Olden & S. J. Ormerod, 2019. Emerging threats and persistent conservation challenges for freshwater biodiversity. Biological Reviews 94(3):849–873.

Rognes, T., T. Flouri, B. Nichols, C. Quince & F. Mahé, 2016. VSEARCH: a versatile open source tool for metagenomics. PeerJ 4:e2584.

Ruffo, S. & F. Stoch, 2006. Checklist and distribution of the Italian fauna. Commune di Verona.

Sari, A., M. Duran & F. Bardakci, 2012. Discrimination of Orthocladiinae species (Diptera: Chironomidae) by using cytochrome c oxidase subunit I. Acta Zoologica Bulgarica 4:73–80.

Schmidt, T. S., J. F. Matias Rodrigues & C. von Mering, 2015. Limits to robustness and reproducibility in the demarcation of operational taxonomic units. Environmental microbiology 17(5):1689–1706.

Schoch, C. L., K. A. Seifert, S. Huhndorf, V. Robert, J. L. Spouge, C. A. Levesque, W. Chen & F. B. Consortium, 2012. Nuclear ribosomal internal transcribed spacer (ITS) region as a universal DNA barcode marker for Fungi. Proceedings of the National Academy of Sciences 109(16):6241–6246.

Schoolmann, G., F. Nitsche & H. Arndt, 2015. Aspects of the life span and phenology of the invasive freshwater shrimp Atyaephyra desmarestii (Millet, 1831) at the northeastern edge of its range (upper Rhine). Crustaceana 88(9):949–962.

Sinclair, C. & S. Gresens, 2008. Discrimination of Cricotopus species (Diptera: Chironomidae) by DNA barcoding. Bulletin of entomological research 98(6):555.

Skoulikidis, N. T., S. Sabater, T. Datry, M. M. Morais, A. Buffagni, G. Dörflinger, S. Zogaris, M. del Mar Sánchez-Montoya, N. Bonada & E. Kalogianni, 2017. Non-perennial Mediterranean rivers in Europe: status, pressures, and challenges for research and management. Science of the Total Environment 577:1–18.

Stewart, K. A., 2019. Understanding the effects of biotic and abiotic factors on sources of aquatic environmental DNA. Biodiversity and Conservation 28(5):983–1001.

Stoch, F., 2000. How many endemic species? Species richness assessment and conservation priorities in Italy. Belgian Journal of Entomology 2(1):125–133.

Strayer, D. L. & D. Dudgeon, 2010. Freshwater biodiversity conservation: recent progress and future challenges. Journal of the North American Benthological Society 29(1):344–358.

Stubbington, R., R. Chadd, N. Cid, Z. Csabai, M. Miliša, M. Morais, A. Munné, P. Pařil, V. Pešić & I. Tziortzis, 2018. Biomonitoring of intermittent rivers and ephemeral streams in Europe: Current practice and priorities to enhance ecological status assessments. Science of the total environment 618:1096–1113.

Taberlet, P., A. Bonin, L. Zinger & E. Coissac, 2018. Environmental DNA: For biodiversity research and monitoring. Oxford University Press.

Tapolczai, K., F. Keck, A. Bouchez, F. Rimet, M. Kahlert & V. Vasselon, 2019. Diatom DNA metabarcoding for biomonitoring: strategies to avoid major taxonomical and bioinformatical biases limiting molecular indices capacities. Frontiers in Ecology and Evolution 7:409.

Tapolczai, K., G. B. Selmeczy, B. Szabó, B. Viktória, F. Keck, A. Bouchez, F. Rimet & J. Padisák, 2021. The potential of exact sequence variants (ESVs) to interpret and assess the impact of agricultural pressure on stream diatom assemblages revealed by DNA metabarcoding. Ecological Indicators 122:107322.

Turon, X., A. Antich, C. Palacín, K. Præbel & O. S. Wangensteen, 2020. From metabarcoding to metaphylogeography: separating the wheat from the chaff. Ecological Applications 30(2):e02036.

Vamos, E., V. Elbrecht & F. Leese, 2017. Short COI markers for freshwater macroinvertebrate metabarcoding. Metabarcoding and Metagenomics, 1, e14625.

Vinarski, M. V., 2017. The history of an invasion: phases of the explosive spread of the physid snail Physella acuta through Europe, Transcaucasia and Central Asia. Biological invasions 19(4):1299–1314.

Vörösmarty, C. J., P. B. McIntyre, M. O. Gessner, D. Dudgeon, A. Prusevich, P. Green, S. Glidden, S. E. Bunn, C. A. Sullivan & C. R. Liermann, 2010. Global threats to human water security and river biodiversity. nature 467(7315):555–561.

Wallace, J. B., J. W. Grubaugh & M. R. Whiles, 1996. Biotic indices and stream ecosystem processes: results from an experimental study. Ecological applications 6(1):140–151.

Weigand, H., A. J. Beermann, F. Čiampor, F. O. Costa, Z. Csabai, S. Duarte, M. F. Geiger, M. Grabowski, F. Rimet & B. Rulik, 2019. DNA barcode reference libraries for the monitoring of aquatic biota in Europe: Gap-analysis and recommendations for future work. Science of the Total Environment 678:499–524.

Westcott, S. L. & P. D. Schloss, 2015. De novo clustering methods outperform reference-based methods for assigning 16S rRNA gene sequences to operational taxonomic units. PeerJ 3:e1487.

Zizka, V. M., V. Elbrecht, J. N. Macher & F. Leese, 2019. Assessing the influence of sample tagging and library preparation on DNA metabarcoding. Molecular ecology resources 19(4):893–899.

Zizka, V. M. A., M. Weiss & F. Leese, 2020. Can metabarcoding resolve intraspecific genetic diversity changes to environmental stressors? A test case using river macrozoobenthos. Metabarcoding and Metagenomics 4:e51925.

Zukowski, S. & K. F. Walker, 2009. Freshwater snails in competition: alien Physa acuta (Physidae) and native Glyptophysa gibbosa (Planorbidae) in the River Murray, South Australia. Marine and Freshwater Research 60(10):999–1005.

