## Supplementary figures and images for "Fresh insights into Mediterranean biodiversity: Environmental DNA reveals spatio-temporal patterns of stream invertebrate communities on Sicily"

### Fig. S1

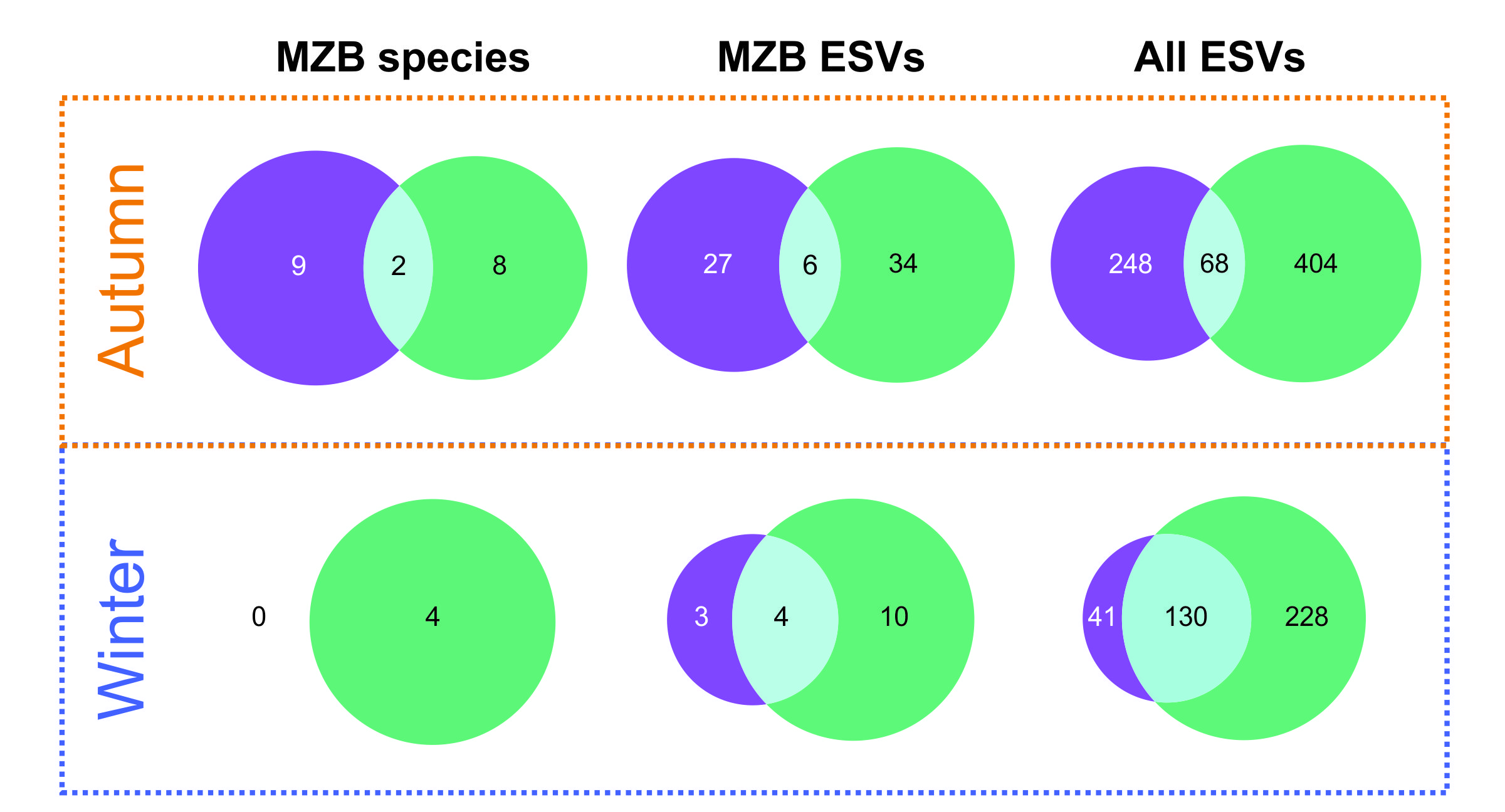

### Fig. S2

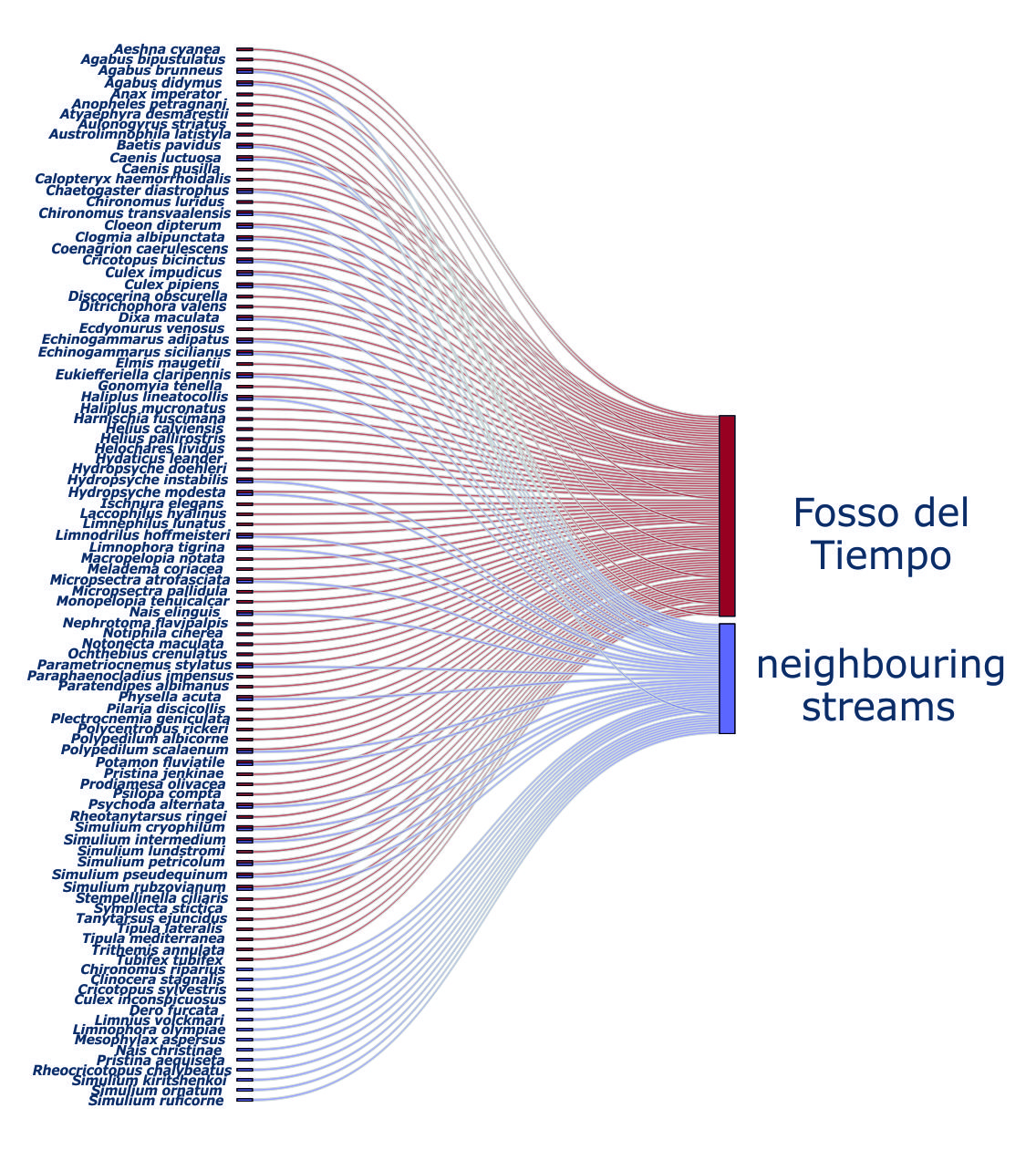
